# Single-cell RNA-seq differential expression tests within a sample should use pseudo-bulk data of pseudo-replicates

**DOI:** 10.1101/2023.03.28.534443

**Authors:** Christoph Hafemeister, Florian Halbritter

## Abstract

Single-cell RNA sequencing (scRNA-seq) has become a standard approach to investigate molecular differences between cell states. Comparisons of bioinformatics methods for the count matrix transformation (normalization) and differential expression (DE) analysis of these data have already highlighted recommendations for effective between-sample comparisons and visualization. Here, we examine two remaining open questions: (i) What are the best combinations of data transformations and statistical test methods, and (ii) how do pseudo-bulk approaches perform in single-sample designs? We evaluated the performance of 343 DE pipelines (combinations of eight types of count matrix transformations and ten statistical tests) on simulated and real-world data, in terms of precision, sensitivity, and false discovery rate. We confirm superior performance of pseudo-bulk approaches without prior transformation. For within-sample comparisons, we advise the use of three pseudo-replicates, and provide a simple R package *DElegate* to facilitate application of this approach.

## 1 Introduction

To learn about the molecular processes that characterize biological conditions, high-throughput profiling of the transcriptome by RNA sequencing (RNA-seq) followed by differential expression (DE) analysis has become a standard workflow in genomics. With the introduction of single-cell RNA-seq (scRNA-seq) technologies this workflow has been expanded to the level of cell types and cell states. As these technologies have seen widespread adoption, the number of methods to perform data normalization, and DE analysis has also grown. However, previous systematic evaluations of DE approaches did not cover the combinations of transformations and DE methods, leaving a knowledge gap we aim to fill here.

The main input to a scRNA-seq analysis is a gene expression count matrix. The elements in this matrix correspond to the number of molecules assigned to a particular gene and cell. For homogeneous cell populations, the vast majority of genes has very low counts (< 1 on average) and follows a Poisson distribution, while highly expressed genes show a variance larger than the mean and can be modeled as negative binomial [Grün et al., 2014, Chen et al., 2018, Hafemeister and Satija, 2019]. However, cell-to-cell variability in gene expression is obscured by technical effects like sequencing depth that may distort the observed counts by several orders of magnitude. Therefore, a considerable amount of effort has gone into developing and examining count matrix transformations (“normalization”) that aim to address the technical variability and mean-variance relationship in the data, with the aim to facilitate downstream analyses like dimensionality reduction and clustering [Hafemeister and Satija, 2019, Lause et al., 2021, Ahlmann-Eltze and Huber, 2023, Booeshaghiet al., 2022, Choudhary and Satija, 2022].

Some commonly used tools for bulk RNA-seq DE analysis have been adopted to work with the sparser input and higher number of observations in scRNA-seq (e.g. single-cell parameterizations of *DESeq2* [Love et al., 2014], and *edgeR* [McCarthy et al., 2012]), and single-cell specific methods were developed (e.g. *MAST* [Finak et al., 2015]). On the other hand, especially with higher cell numbers, general-purpose statistical tests, applied independently per gene, have been used instead (e.g., randomization, Wilcoxon rank-sum test, t-test, logistic regression). As an alternative to treating each cell as an observation in the DE test, bulk RNA-seq DE methods may be used after aggregating single-cell counts (e.g., original use-case of *DESeq2, edgeR, limma* [Law et al., 2014]), however, this would normally require multiple samples.

Initial comparisons of tools for single-cell DE analysis highlighted inconsistencies between methods and found that tools designed specifically for single-cell data did not necessarily outperform general-purpose methods [Soneson and Robinson, 2018]. Later benchmarks included methods that aggregate single cells, or use multi-sample designs [Pullin and McCarthy, 2022, Crowell et al., 2020, Squair et al., 2021]. Jointly, their results highlighted the importance of biological replicates and methods that account for them, and demonstrated the effectiveness of aggregating data (by creating pseudo-bulk) for DE analysis.

Here, we aimed to expand previous benchmarks of single-cell DE methods by focusing on single-sample designs and considering recently developed data transformations. We wanted to address two questions:

1. What are best combinations of data transformations and statistical tests for single-cell DE analysis?
2. How do pseudo-bulk approaches perform in the absence of real replicates?

We approached the task of finding DE genes as a sequence of a data transformation step and a statistical test. In total, we combined parameterizations of eight types of count matrix transformations and ten test methods for a total of 343 DE “pipelines” (Fig. 1, Supp. Tables 1-4). We focused on comparing two groups of cells in the absence of replicates, as is commonly done when working within an individual sample (e.g., comparing cell types or cell states from one individual), and evaluated each pipeline on simulated data using precision, sensitivity, Matthews correlation coefficient, and false discovery rate. Additionally, we performed evaluation using real-word data with independent validation data.

**Figure 1:**
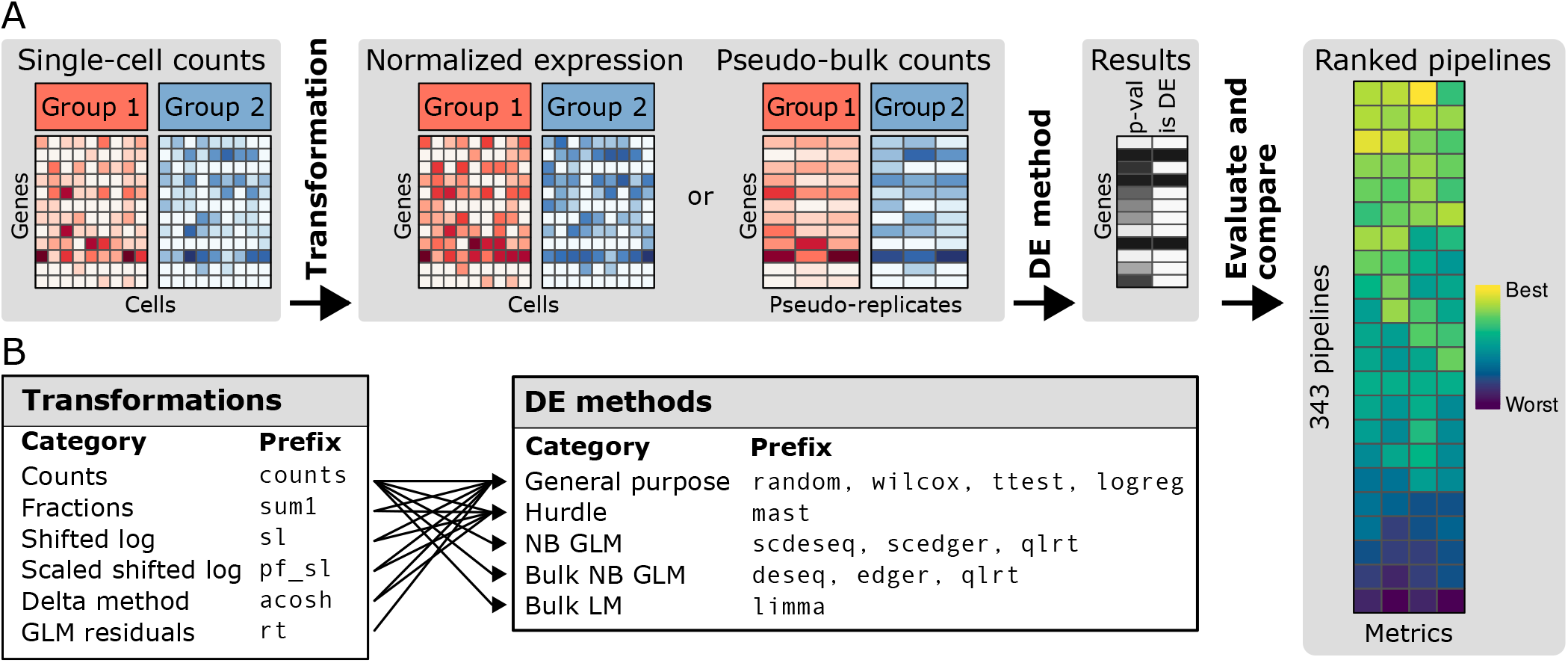
Evaluation of an exhaustive collection of DE pipelines. A) In our terminology, each pipeline is a combination of one particular data transformation method and one particular statistical test. All pipelines were evaluated jointly on simulated data and an immune data set. B) Transformations were paired with DE methods. We use a naming scheme with unique prefixes to indicate the methods. NB negative binomial, GLM generalized linear model, LM linear model.

## 2 Results

Overall we tested 343 DE pipelines (combinations of commonly used count-data transformations and statistical tests) on four types of simulation experiments, and on one immune cell dataset.

### 2.1 Simulated data: Pseudo-bulk methods with pseudo-replicates have lower FDR and outperform single-cell methods

First, we sought to evaluate the DE pipelines in well-controlled settings, in which we knew beforehand which genes were truly differentially expressed and which were not. Therefore, we simulated an artificial dataset of four distinct experiment types using the *muscat* package [Crowell et al., 2020] (Fig. 2A, for details see Methods). In each experiment, we simulated two groups. Three experiments covered balanced cell numbers (20 vs. 20, 200 vs. 200, 2k vs. 2k), while one represented an unbalanced comparison (200 vs. 2k). In all cases, we introduced a fixed ratio of 10% DE genes, and generated 13 replicates.

**Figure 2:**
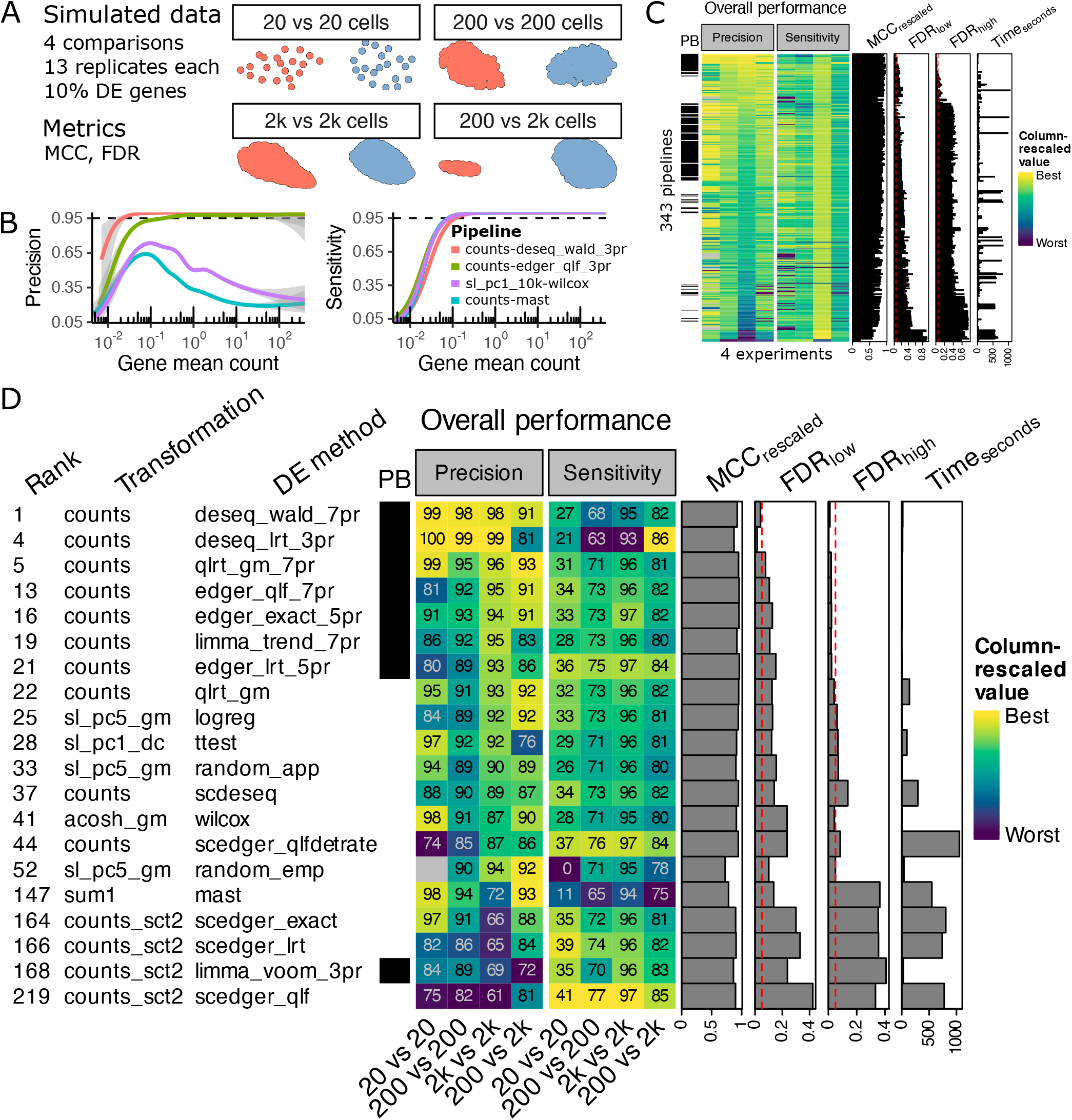
Simulation experiments. A) We performed four types of simulation experiments with varying cell numbers. B) Example results for four pipelines, evaluating precision and sensitivity as function of mean expression between groups in the 2k vs. 2k cell experiment. C) Overview of evaluation results. 343 DE pipelines (combination of transformation and test method) were ranked based on Matthews correlation coefficient (MCC), and false discovery rate (FDR) among lowly or highly expressed genes. D) Top-performing pipelines for every test method in the benchmark. For every test method, only the highest-ranking transformation is shown. A black entry in the PB column indicates pipelines using pseudo-bulk data. The numbers in the heatmap show precision and sensitivity as percent for the four experiments performed. For the exact ranking procedure see the Methods section.

For every experiment, replicate, and pipeline we determined the predicted DE genes (at an expected FDR cutoff of 0.05 based on the method’s own calculated P-values) and calculated precision, sensitivity, and Matthews correlation coefficient (MCC, re-scaled per experiment, see Methods section) across all genes. Additionally, we evaluated highly and lowly expressed genes separately and calculated the true FDR, averaging across replicates. The decision to separately measure FDR for genes based on mean expression was motivated by prior reports of high FDR for highly expressed genes for some methods [Squair et al., 2021] and our own observations of the mean vs. precision dependency (examples shown in Fig. 2B). We aimed to compile a summary that captured overall performance in all experiments and that highlighted methods that did well across all expression levels. Therefore, we combined the MCC, a measure of the overall agreement of the predicted DE genes with the ground truth, with the true FDR for lowly and highly expressed genes. Especially for highly expressed genes, FDR can be a problem that would bot be captured by a genome-wide metric due to the low number of genes affected. The final aggregate score is then MCC+(1–FDR_low_)+(1–FDR_high_), where the MCC is the average of the rescaled MCC values for the four experiments. The FDR values are taken from the experiment with the highest number of cells, as that experiment tended to yield more false positives. See Fig. 2C and Additional Files 1&2 for complete results. The highest ranking pipeline for every DE method is shown in Fig. 2D.

We found that the top-ranking pipelines all employed test methods designed to work on raw counts from multiple replicates of bulk data. In our single-sample setting, these tests had been run using pseudo-bulk data from random splits of the dataset (“pseudo-replicates”, see Methods section) without any further count data transformations applied. These were also the methods that most effectively controlled the true FDR among the highly expressed genes. Specifically, the highest-ranking methods were *DE-seq2, qlrt, edgeR,* and *limma trend.* While all four methods had high sensitivity when many cells were compared, we found that *DESeq2* had the highest precision at the cost of slightly lower sensitivity. This conservative DE calling made *DESeq2* particularly good at controlling the FDR for lowly expressed genes (mean ≤ 0.1). Methods directly using the single-cell data ranked in the middle, with tests specifically modeling count data (*qlrt*, and single-cell-adapted *DESeq2* and *edgeR*) performing comparably to generic statistical tests. The latter were paired with prior scaling transformations (e.g., logistic regression with log-transformed and median-centered counts), and reported fewer false positives among the highly expressed genes. The bottom of the ranking was made up of methods that clearly failed to control the overall FDR, especially with an increasing number of cells tested. These methods have in common that they were designed to take read counts as input, but did not perform well with the tested parameter choices. In particular, an inappropriate choice of size factor models led to low precision (e.g., *qlrt* which worked well with raw counts did not perform well with some size factors). Notably, a closer look at the full results table revealed that the popular combination of log-counts per 10k transformation and Wilcoxon rank sum test was among those low-ranking pipelines (position 198/343, Add. File 1).

### 2.2 Immune cell data: Pseudo-bulk methods have highest agreement with independent bulk data results

To investigate the performance of the DE pipelines on real data, we contrasted the three main immune cell compartments (monocytes, B cells, T cells) from a commonly used dataset of peripheral blood mononuclear cells from a single healthy donor [10x Genomics]. We used bulk RNA-seq from sorted cell populations to determine a high-confidence set of DE genes that we considered a ground truth reference. In short, we queried the BLUEPRINT consortium data (http://dcc.blueprint-epigenome.eu) for RNA-seq experiments of venous blood samples for cell types that match our three main cell types. This resulted in 9 B-cell, 10 monocyte, and 17 T-cell samples. Bulk data DE genes were then determined by running *edgeR, limma,* and *DESeq2,* and combining the results by intersection (Fig. 3A).

**Figure 3:**
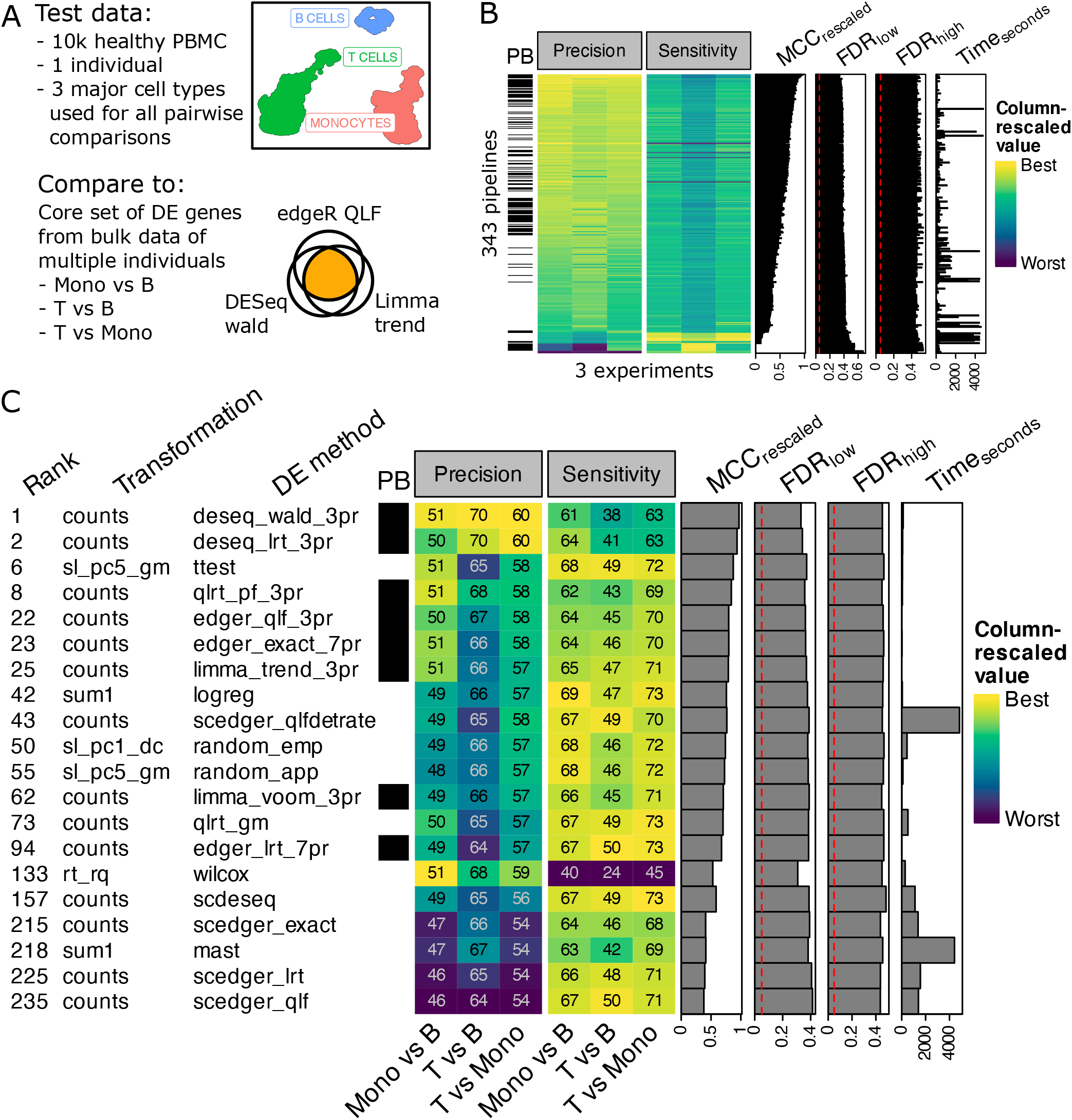
Immune cell experiments. A) A single-cell dataset of healthy PBMC was used for testing the pipelines. For evaluation, a dataset of cell-type-sorted bulk RNA-seq was used. B) All pipelines were evaluated and ranked according to relative performance. C) Top-performing pipelines for every test method (as in Fig. 2).

All DE pipelines were evaluated and ranked using the same metrics as for the simulated data, with the exception that FDR values shown are averages across all three comparisons (Fig. 3B). Again, among the high-ranking methods we saw mostly bulk data methods with raw counts as input *(DESeq2, qlrt, edgeR, limma trend),* and a t-test pipeline as a notable exception (Fig. 3 and Add. Files 3&4). The FDR values observed in these experiments are generally much higher than in the simulation experiments with the smallest value being around 0.3 for the lowly expressed genes, and 0.4 for the highly expressed genes. This means that all pipelines returned a high proportion of DE genes that are not in the multi-sample derived ground truth.

### 2.3 Well-established bulk methods using untransformed pseudo-bulk counts consistently outperform other pipelines

To establish what pipelines consistently performed well in our tests, we ordered the pipelines by their worst rank across the two evaluation scenarios (simulated data, immune cell data) and focused on the pipelines that ranked in the top 50 in both cases (Fig. 4). Among these 23 top-performers, 15 are using the pseudo-replicate pseudo-bulk approach with untransformed counts, and nine of these make up the top of the list (*DESeq2, qlrt, edgeR,* and *limma trend* are all represented). The single-cell methods in this list are t-test (5 times), logistic regression (2 times), and a single-cell optimized variant of *edgeR*. Regarding the count data transformations, we observed that in six out of the eight cases the single-cell pipelines used a simple shifted-log with cell size factors defined by deconvolution (*scran*) or the geometric mean (see Supplementary Table 2 for size factor details).

**Figure 4:**
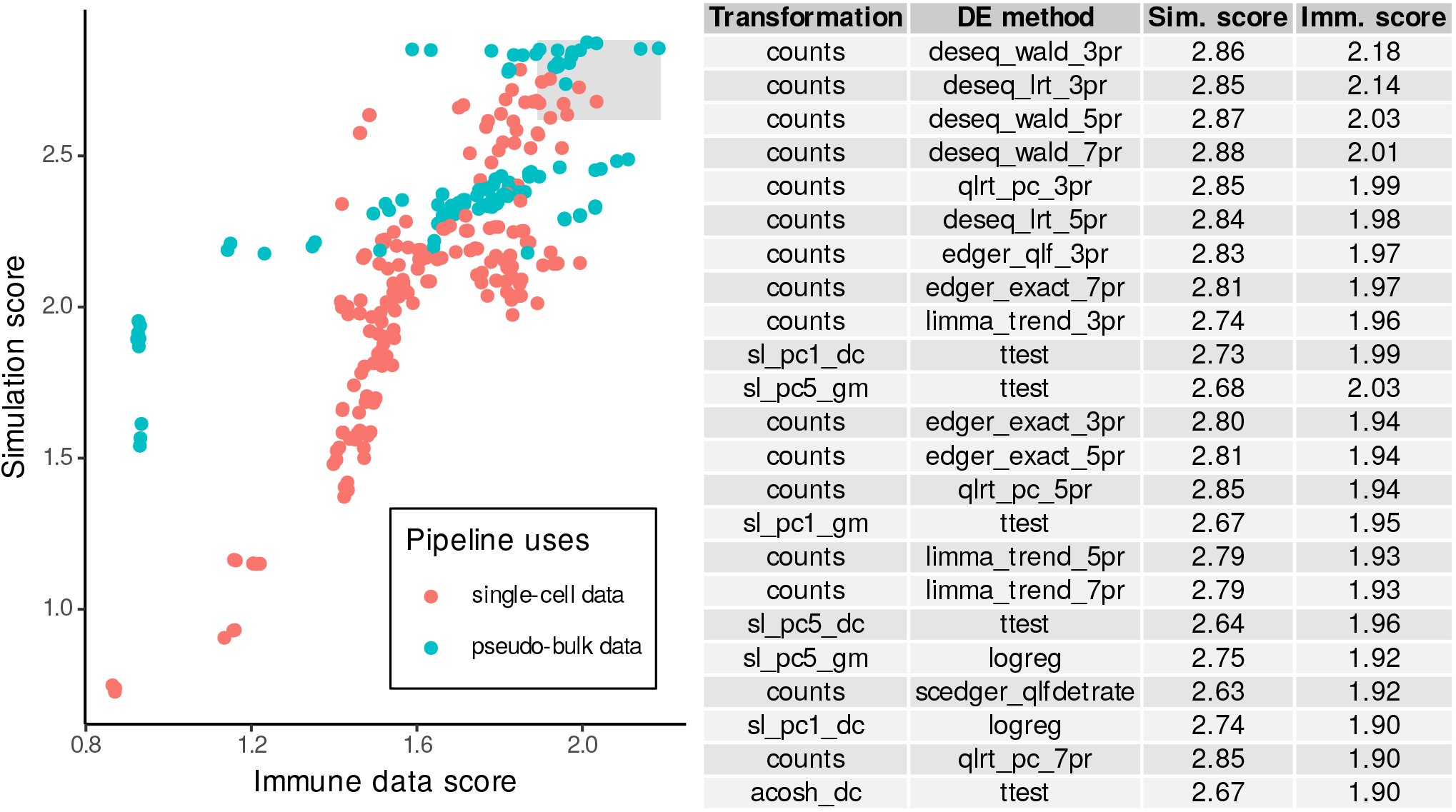
Simulation and immune data results combined. We plot all pipelines with their immune cell experiment score and their simulation data score. The table and the gray rectangle show all pipelines that are ranked in the top 50 of both experiment types. Pipelines are ordered by their worst rank.

Given that pipelines using pseudo-replicate (PR) data consistently ranked highly in our benchmark, we sought to examine the influence of the choice of number of PRs on the result. In all experiments, we had included runs with 3, 5, or 7 PRs, to evaluate the impact of that parameter. We averaged the simulation and immune cell evaluation scores and assessed performance as a function of the number of PRs (Supp. Fig. 1). We found that three PRs mostly led to better performance than a higher number of PRs (true for *qlrt, limma voom, edgeR QLF, DESeq2).* In a few cases, three PRs resulted in only slightly lower, or very similar performance than more PRs *(limma trend, edgeR LRT, edgeR exact*).

### 2.4 Pipeline runtimes

While we did not focus on pipeline runtimes for evaluation, we kept track of the time it took to perform transformation and DE testing (Supp. Fig. 2&3). Transformations that required model parameter estimation were among the slowest ones, as were those that used *scran*’s deconvolution-based cell size factor. Among the DE tests applied to single-cell data, highly efficient implementations of the t-test, logistic regression, and Wilcoxon rank-sum test stood out as particularly fast, while all model-based approaches were significantly slower (at least 50x). Methods applied to aggregated counts were also fast, with *limma* and *edgeR* being the fastest.

## 3 Discussion

In this study, we set out to evaluate the performance of differential expression (DE) pipelines on simulated and real single-cell RNA-seq data. We included many combinations of data transformations and statistical tests, as well as test methods originally designed for bulk RNA-seq. Since previous studies have already convincingly shown that bulk methods are the methods of choice when replicates are given [Crowell et al., 2020, Squair et al., 2021], we focused on the scenario where no replicates are available. For example, this would be the case when trying to find markers of different clusters in one sample.

The results of our simulation experiments showed that bulk methods that use pseudo-bulk raw count data from pseudo-replicates ranked highest and were most effective in controlling the false discovery rate (FDR) for highly expressed genes. This is in line with previous observations [Squair et al., 2021]. For real scRNA-seq data, the top-performing pipelines were also dominated by the same kind of pipelines, but the differences between single-cell and pseudo-replicate methods were less clear. This is in part due to the different nature of the evaluation – biological replicates were used to generate the set of true DE genes, while the evaluation was done using a single sample resulting in all pipelines suffering from higher false positives. Another difference is the smaller variability in sequencing depth between cell groups being compared – in the simulation experiments one group was downsampled to 85%, while the differences in sequencing depth within the singlecell immune data were very small. This decreased the effect of cell size factors and library size normalization on the overall outcome.

Among the top-performing single-cell pipelines, we found mostly shifted-log transformed count data in combination with a t-test or logistic regression. This is an interesting result, showing that rather simple transformations, when using the right size factors (here geometric mean, or *scran*’s size factors), can be appropriate for more generalized statistical analyses. This finding matches the good performance Ahlmann-Eltze and Huber [2023] have recently reported for the shifted-log transformation in the context of dimensionality reduction, and the highlighting of t-test, logistic regression, and Wilcoxon rank-sum test by Pullin and McCarthy [2022] for marker gene identification.

Overall, our results indicate that DE pipelines designed for bulk sequencing data are the method of choice also in the single-cell single-replicate scenario. We recommend the use of pseudo-bulk data of three pseudo-replicates, and *DESeq2* (lowest FDR) with Wald test, or *edgeR* (slightly higher sensitivity) with QLF test. Besides being more accurate, these pipelines are also among the fastest that we tested.

In a more general context of single-cell RNA-seq data analysis, our results support the idea that there is not just one data transformation that is suited for all kinds of tasks. Rather, one should use different transformations for different purposes. For example, shifted-log or Pearson residuals would be better suited for dimensionality reduction [Ahlmann-Eltze and Huber, 2023], mapping to manifolds, and nearest-neighbor network creation, while pseudo-bulk of untransformed counts and pseudo-replicates would be better for DE testing. The overall favorable results of pseudo-replicate approaches can also be seen as an endorsement of meta-cell methods, as pioneered by Baran et al. [2019], where homogeneous single-cell profiles are grouped for robust statistics (see also Bilous et al. [2022], Persad et al. [2023]).

Our benchmarking study has limitations. For one, we focused on just one particular sample type, namely PBMCs. Using additional samples from other tissues, especially for real-data experiments, could change the pipeline rankings. However, the overall agreement between simulation data results and real data results is an indicator, that we captured true performance differences. Also, the single-cell data used in this pipeline benchmark was generated with 10x Chromium technology and we did not evaluate how the pipelines performed on data generated using other singlecell technologies. Furthermore, for evaluation, we have not considered true or predicted effect size, only statistical significance. This was a deliberate decision, as effect size (e.g., fold change) depends on both, the transformation and DE method, and is therefore not comparable between pipelines. Thus, by focusing on the binary DE call and its effect direction, we simplified the evaluation and could avoid the use of arbitrary thresholds. Finally, we would like to note, that pseudo-replicates are not a substitute for true biological replicates. Only true biological replicates can supply the estimates of inter-individual gene expression variability needed to make robust inferences of DE.

### 3.1 Conclusion

In the benchmark presented here, we see clear evidence for the good performance of bulk DE methods in the single-cell scenario, even in the absence of true replicates. After testing many data transformations and DE tests, we do not see any advantages of true single-cell DE pipelines, leading us to suggest the use of pseudo-bulk data of pseudo-replicates with bulk DE methods *(DESeq2, edgeR)* instead. To facilitate the use of this recommended methodology, we have implemented these methods together with the generation of pseudo-replicates from count matrices, SingleCellExperiment, and Seurat objects in a simple R package *DElegate*.

## 4 Methods

### 4.1 Data

#### 4.1.1 Simulated data

To simulate single-cell data with known DE genes, we used *muscat* [Crowell et al., 2020]. While simulated data may not accurately reflect real data (see Crowell et al. [2022] for a recent discussion), we expect the relative ranking of pipelines to still be informative in this simple two-group experimental design. As an input to *muscat*, we used two samples: 1) the monocyte cluster from the single-cell immune data described below, and 2) the same cells, but each cell downsampled to 85% of the total counts. We downsampled one sample, to make the task of DE testing more challenging and more similar to real data situations, where total counts may vary between cell types, or may arise from sequencing depth differences. For every simulation experiment, we generated 13 replicates of two samples with 10% genes DE (using the DE *muscat* model only).

#### 4.1.2 Immune cell data

As “real” single-cell RNA-seq test data, we used a publicly available dataset of approximately 10k healthy human PBMC [10x Genomics]. We then performed filtering to keep only high-quality cells: 1. We excluded cells with more than 15% mitochondrial reads. 2. We z-scored the log10-transformed number of genes detected and excluded cells with values outside the range [−3, 3]. 3. We z-scored the log10 of the number of reads and excluded cells outside of [−3, 3]. We then used *Seurat* v4.1.0 [Hao et al., 2021] to cluster the cells (we used the following non-default parameters: PCs = 15, distance metric = manhattan, k parameter = 20, n neighbors = 40, cluster resolution = 0.8). We then used *celltypist* [Domínguez Conde et al., 2022] with the Immune_All_Low model and the best_match option to annotate the cells. To focus on the three main immune cell types, we merged the subtypes of T cells, monocytes, and B cells, respectively. Finally, we kept all clusters that had not more than 50% “other” cells (e.g., NK cells) and that were not less than 100 cells in size, and removed all “other” cells. This resulted in a data set of 4,493 T cells, 4,453 monocytes, and 1,028 B cells.

The immune cell validation data was obtained from the BLUEPRINT consortium data site at http://dcc.bueprint-epigenome.eu. Using the provided metadata, we selected RNA-seq experiments of venous blood samples for cell types that match our three main immune cell types described above. This resulted in 9 B-cell samples from 6 individuals, 10 monocyte samples from 8 individuals, and 17 T-cell samples from 12 individuals. Bulk data DE genes were then determined by running three bulk DE methods (*edgeR QLF*, *limma trend*, and *DESeq2 Wald*) with default parameters, and combining the results by intersection (FDR ≤ 0.05 per method).

### 4.2 Evaluation metrics and pipeline ranking

#### 4.2.1 Simulation experiments

In all simulation experiments, “positives” were defined as the genes identified as DE by a given pipeline (at FDR cutoff of 0.05). True positives (TP) are therefore those positives that were simulated with a non-zero fold change, while false positives (FP) were simulated with a fold change of 0. True negatives (TN) are genes not identified as DE and simulated with a fold change of 0, while false negatives (FN) were simulated with a non-zero fold change. In the event of positives with wrong effect direction (e.g., simulated with positive fold change, but identified as DE with negative fold change), the gene is counted as 1/2 FP and 1/2 FN.

For every experiment we calculate the following metrics:

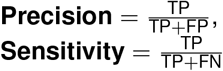

##### Matthews correlation coefficient (MCC)

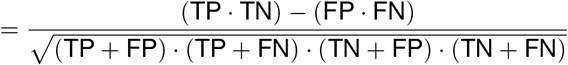

We clipped the MCC to a lower bound of 0 to have a wider dynamic range for the better-than-random results (MCC below 0 indicates a bias towards misclassification).

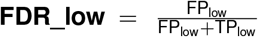. The FDR among all genes with a geometric mean <0.1 across the simulation group means. As lowly and highly expressed genes follow different distributions [Grün et al., 2014, Chen et al., 2018, Hafemeister and Satija, 2019], it was important to consider both groups of genes separately to evaluate DE pipelines.

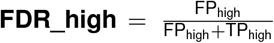. The FDR among all genes with a geometric mean ≥ 1 across the simulation group means.

##### Final ranking

To rank the pipelines, we calculated a score for each pipeline: MCC_rescaled_ + (1 - FDR_low_) + (1 - FDR_high_). With this score we aimed to capture the overall agreement of the predicted set of DE genes with the ground truth, but also penalized pipeline biased towards false positives among lowly and highly expressed genes. In detail, we rescaled MCC to range from 0 to 1 across all pipelines per experimental replicate, then averaged (mean) the replicates per experiment and pipeline, and then averaged (mean) the experiments to obtain one MCCrescaled score per pipeline. For the FDR, we used the 2k vs. 2k cell experiment and averaged the values across replicates.

#### 4.2.2 Immune cell data experiments

For the evaluation of the DE pipelines on real data, we performed DE analysis on bulk RNA-seq data (see Data section above), and treated the results as a proxy for a real “ground truth” (high-confidence set of DE genes). Specifically, for every cell type comparison, we used the intersection of the DE genes by 1) *edgeR* with QLF test, 2) *DESeq* with Wald test, 3) *limma-trend.* For every method we used an estimated FDR ≤ 0.05, and we required all three DE methods to also agree on the effect direction to add a gene to the DE set.

The definition of the pipeline evaluation metrics was then analogous to the simulation experiments. Note that we expected lower precision and higher FDR than compared to the simulation experiments. Our ground truth represented a core set of DE genes derived from bulk data with replicates from multiple individuals, while the test data consists of a single sample from one individual, assayed with a different technology. However, all pipelines should be affected by these differences, and the metrics still allowed us to rank the pipelines according to their overall agreement with the bulk results.

##### Final ranking

Similarly to the simulation experiments, we calculated a score for each pipeline: MCC_rescaled_ + (1 - FDR_low_) + (1 - FDR_high_). We rescale MCC to range from 0 to 1 across all pipelines per experiment, and then averaged the experiments to obtain one MCCrescaled score per pipeline. For the FDR, we averaged the values across experiments.

## Additional files

Additional File 1: CSV file with details of simulation experiment results

Additional File 2: PDF file with heatmap of simulation experiment results

Additional File 3: CSV file with details of immune cell experiment results

Additional File 4: PDF file with heatmap of immune cell experiment results

## Data and software availability

We have uploaded the data used in our experiments to zenodo where it is publicly available at https://doi.org/10.5281/zenodo.7751830.

All code used to execute the benchmark, and a project Dockerfile are available on GitHub at https://github.com/cancerbits/scDE_benchmark.

The *DElegate* R package facilitating the use of recommended pipelines is available at https://github.com/cancerbits/DElegate.

## Author contributions

Conceptualization: C.H., F.H. Formal Analysis: C.H. Investigation: C.H. Methodology: C.H. Project administration: F.H. Resources: F.H. Software: C.H. Supervision: F.H. Visualization: C.H. Writing – original draft: C.H., F.H. Writing – review & editing: C.H., F.H.

## Acknowledgements

This study makes use of data generated by the BLUEPRINT Consortium. A full list of the investigators who contributed to the generation of the data is available from https://www.blueprint-epigenome.eu. Funding for the BLUEPRINT project was provided by the European Union’s Seventh Framework Programme (FP7/2007-2013) under grant agreement no 282510.

## Supplementary tables

**Table 1:** Transformations / normalizations used

| Transformation | Abbreviation | Details | Reference |
| --- | --- | --- | --- |
| Raw counts | counts | No transformation - raw count matrix used as input | NA |
| Fractions | sum1 | Counts of each cell are scaled to a total of 1 | NA |
| Shifted log | sl_pc(1\|5),<br>sl_pc(1\|5)_{sf} | Log transformation after scaling each cell by a size factor ({sf} is replaced with size factor abbreviation) and adding a pseudo-count of 1 or 5 | several, see size factor table below |
| Scaled shifted log | pf_sl_pf | Like sl_pc1_pf above but followed by another round of cell-scaling using proportional fitting (pf) | Booeshaghi et al. [2022] |
| Delta method | acosh,<br>acosh_{sf} | Variance-stabilizing transformation for Negative Binomial (Gamma-Poisson) data as implemented in transformGamPoi::acosh_transform(); cells are scaled using various methods to calculate cell size factors and a default overdispersion of alpha = 0.05 is used | Ahlmann-Eltze and Huber [2023] |
| GLM residuals | rt_pr_sct,<br>rt_pr_sct2,<br>rt_pr_tgp,<br>rt_rq | Fit a generalized linear model for Negative Binomial count data that corrects for sequencing depth and return the Pearson residuals. rt_pr_sct and rt_pr_sct2 are implemented in sctransform::vst() without and with setting vst.flavor = 'v2'. rt_pr_tgp is similar to rt_pr_sct above but using the implementation in transformGamPoi::residual_transform() and run with default parameters (e.g., no clipping). rt_rq is like rt_pr_tgp above, but returns quantile-randomized residuals. Quantile-randomized residuals match the cumulative density function of the Negative Binomial (Gamma-Poisson) and the Normal distribution. Implemented in transformGamPoi::residual_transform() and run with default parameters | Hafemeister and Satija [2019], Choudhary and Satija [2022], Ahlmann-Eltze and Huber [2023] |
| GLM-corrected counts | counts_sct,<br>counts_sct2 | Sequencing-depth-corrected counts. An sctransform model is fit and the regression model reversed keeping the residuals fixed and assuming the same total count per cell. The corrected counts are rounded to the nearest integer and negative values clipped to 0. Implemented in sctransform::correct_counts() | Hafemeister and Satija [2019], Choudhary and Satija [2022] |
| Shifted log of corrected counts | sl_pc1_sct,<br>sl_pc1_sct2 | Like above but after correcting the counts a pseudo-count of 1 is added and log-transformation is applied | Hafemeister and Satija [2019], Choudhary and Satija [2022] |

**Table 2:** Single-cell size factors used

| Abbreviation | Details | Reference |
| --- | --- | --- |
| 10k | Scale each cell to a total of 10,000 | e.g., Satija et al. [2015], Wolf et al. [2018] |
| pf | Proportional fitting; scale each cell to a total that is the mean of the totals of all cells | Booeshaghi et al. [2022] |
| ns | Normed sum; scale each cell by a size factor that is proportional to the total counts; size factors are scaled to have geometric mean of 1 | Ahlmann-Eltze and Huber [2023] |
| pc | Poscounts; modification of the median ratio method introduced in DESeq; implemented in transformGamPoi package; size factors are scaled to have geometric mean of 1 | Anders and Huber [2010], Ahlmann-Eltze and Huber [2023] |
| dc | Deconvolution; scran's method of deconvolving size factors from cell pools (scran::calculateSumFactors()); size factors are scaled to have geometric mean of 1 | Lun et al. [2016] |
| gm | Geometric mean; cell size factors are proportional to the geometric mean across all genes; to avoid log of zero, we add a small value to all counts; this value is equal to the median detection rate across all cells; size factors are scaled to have geometric mean of 1 | NA |

**Table 3:**
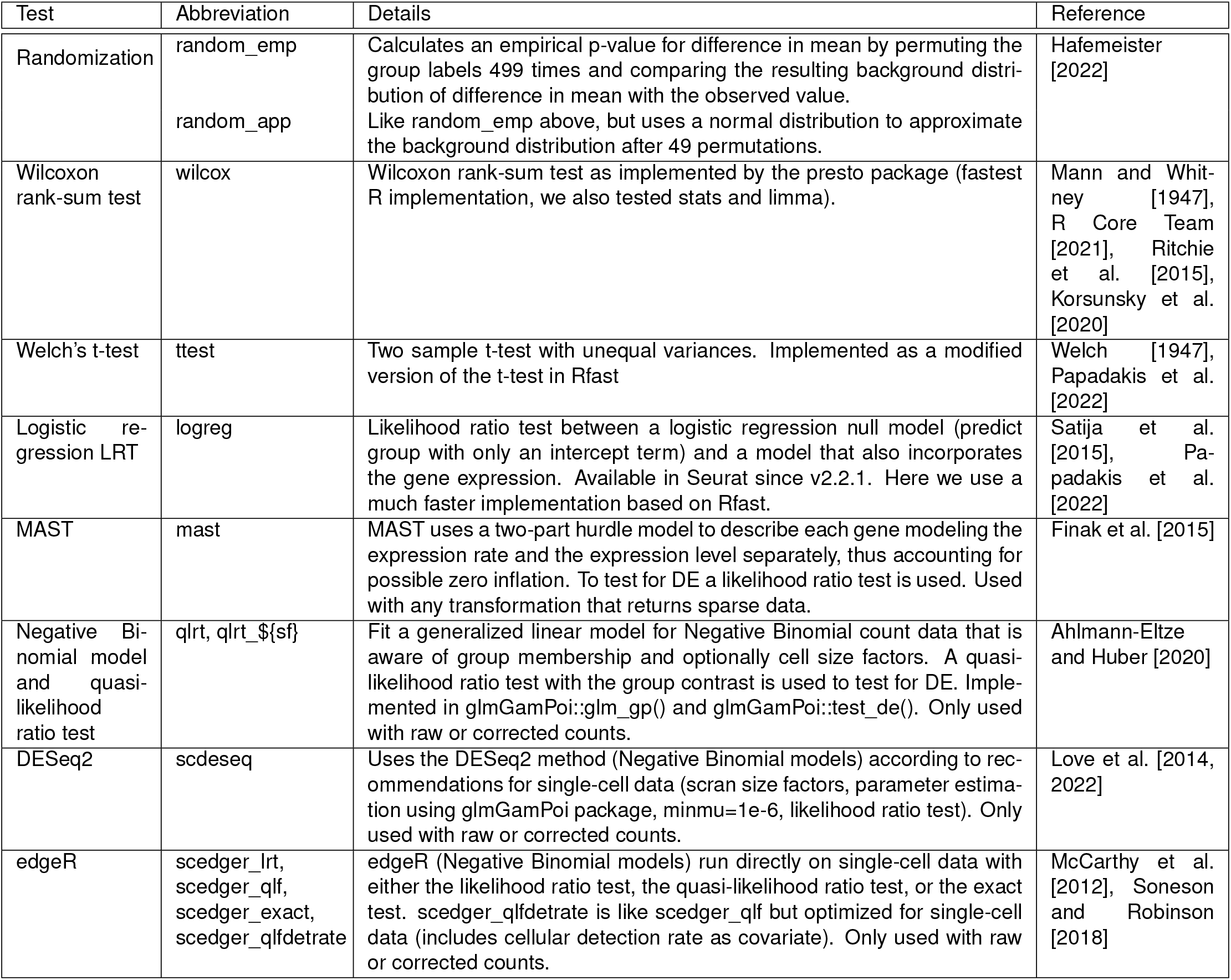
Single-cell differential expression tests used

| Test | Abbreviation | Details | Reference |
| --- | --- | --- | --- |
| Randomization | random_emp<br>random_app | Calculates an empirical p-value for difference in mean by permuting the group labels 499 times and comparing the resulting background distribution of difference in mean with the observed value.<br>Like random_emp above, but uses a normal distribution to approximate the background distribution after 49 permutations. | Hafemeister [2022] |
| Wilcoxon rank-sum test | wilcox | Wilcoxon rank-sum test as implemented by the presto package (fastest R implementation, we also tested stats and limma). | Mann and Whitney [1947], R Core Team [2021], Ritchie et al. [2015], Korsunsky et al. [2020] |
| Welch's t-test | ttest | Two sample t-test with unequal variances. Implemented as a modified version of the t-test in Rfast | Welch [1947], Papadakis et al. [2022] |
| Logistic regression LRT | logreg | Likelihood ratio test between a logistic regression null model (predict group with only an intercept term) and a model that also incorporates the gene expression. Available in Seurat since v2.2.1. Here we use a much faster implementation based on Rfast. | Satija et al. [2015], Papadakis et al. [2022] |
| MAST | mast | MAST uses a two-part hurdle model to describe each gene modeling the expression rate and the expression level separately, thus accounting for possible zero inflation. To test for DE a likelihood ratio test is used. Used with any transformation that returns sparse data. | Finak et al. [2015] |
| Negative Binomial model and quasi-likelihood ratio test | qlrt, qlrt_{\$sf} | Fit a generalized linear model for Negative Binomial count data that is aware of group membership and optionally cell size factors. A quasi-likelihood ratio test with the group contrast is used to test for DE. Implemented in glmGamPoi::glm_gp() and glmGamPoi::test_de(). Only used with raw or corrected counts. | Ahlmann-Eltze and Huber [2020] |
| DESeq2 | scdeseq | Uses the DESeq2 method (Negative Binomial models) according to recommendations for single-cell data (scrn size factors, parameter estimation using glmGamPoi package, minmu=1e-6, likelihood ratio test). Only used with raw or corrected counts. | Love et al. [2014, 2022] |
| edgeR | scedger_lrt,<br>scedger_qlf,<br>scedger_exact,<br>scedger_qlfdetrade | edgeR (Negative Binomial models) run directly on single-cell data with either the likelihood ratio test, the quasi-likelihood ratio test, or the exact test. scedger_qlfdetrade is like scedger_qlf but optimized for single-cell data (includes cellular detection rate as covariate). Only used with raw or corrected counts. | McCarthy et al. [2012], Sonesson and Robinson [2018] |

**Table 4:**
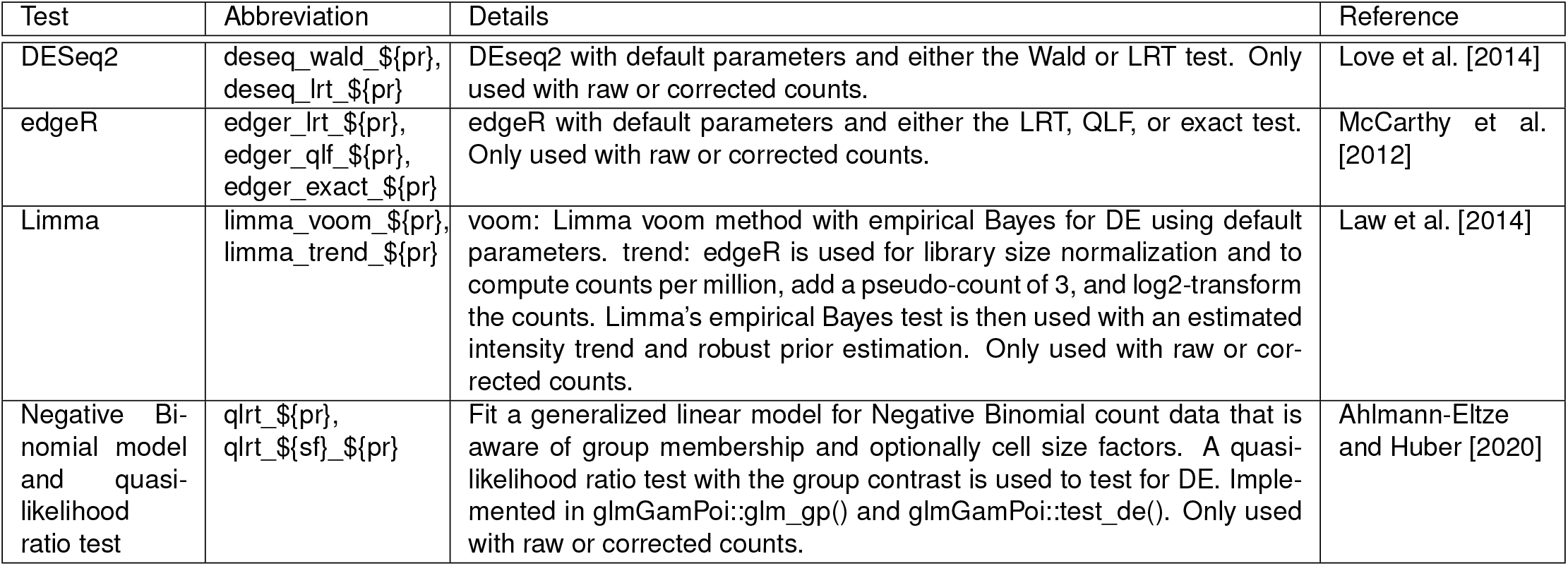
Bulk differential expression tests used

| Test | Abbreviation | Details | Reference |
| --- | --- | --- | --- |
| DESeq2 | deseq_wald_{\$pr},<br>deseq_lrt_{\$pr} | DESeq2 with default parameters and either the Wald or LRT test. Only used with raw or corrected counts. | Love et al. [2014] |
| edgeR | edgeR_lrt_{\$pr},<br>edgeR_qlf_{\$pr},<br>edgeR_exact_{\$pr} | edgeR with default parameters and either the LRT, QLF, or exact test. Only used with raw or corrected counts. | McCarthy et al. [2012] |
| Limma | limma_voom_{\$pr},<br>limma_trend_{\$pr} | voom: Limma voom method with empirical Bayes for DE using default parameters. trend: edgeR is used for library size normalization and to compute counts per million, add a pseudo-count of 3, and log2-transform the counts. Limma's empirical Bayes test is then used with an estimated intensity trend and robust prior estimation. Only used with raw or corrected counts. | Law et al. [2014] |
| Negative Binomial model and quasi-likelihood ratio test | qlrt_{\$pr},<br>qlrt_{\$sf}_{\$pr} | Fit a generalized linear model for Negative Binomial count data that is aware of group membership and optionally cell size factors. A quasi-likelihood ratio test with the group contrast is used to test for DE. Implemented in glmGamPoi::glm_gp() and glmGamPoi::test_de(). Only used with raw or corrected counts. | Ahlmann-Eltze and Huber [2020] |

## Supplementary Figures

**Supp. Figure 1:**
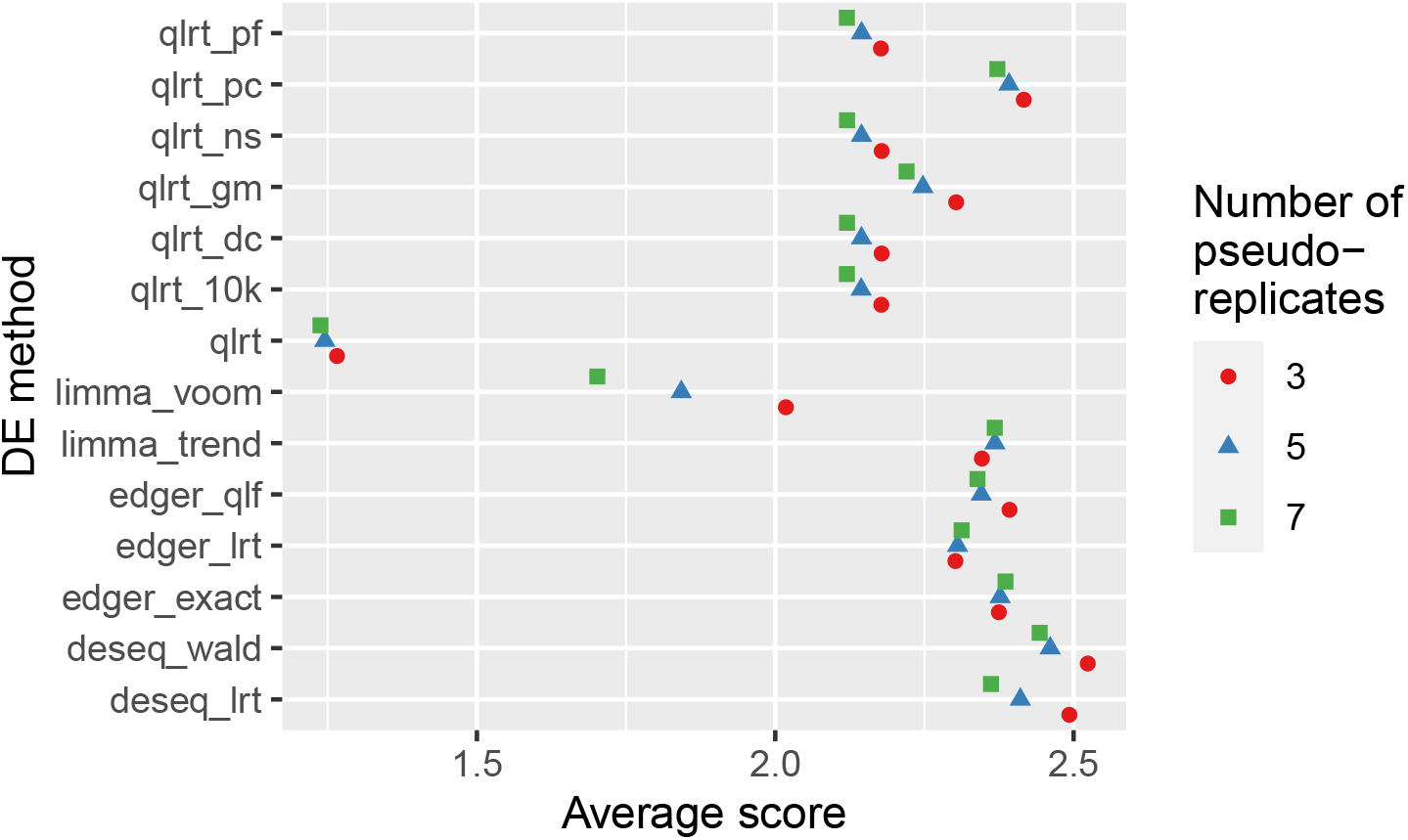
Average score of simulation experiments and immune cell experiments for the different number of pseudo-replicates. Only pipelines with untransformed counts as input are shown.

**Supp. Figure 2:**
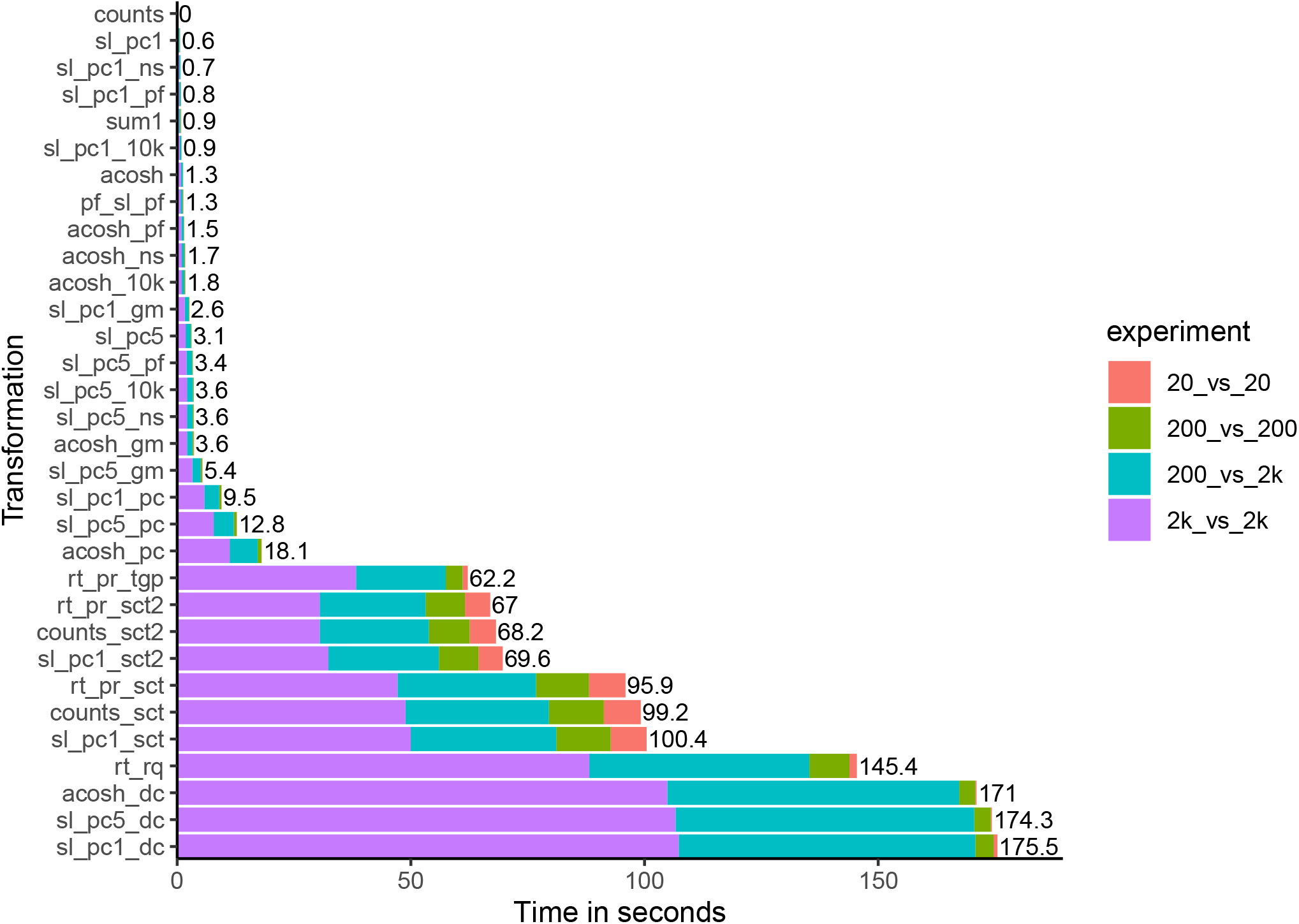
Timing for the different transformations. For each simulation experiment, the average across pipelines and replicates is shown.

**Supp. Figure 3:**
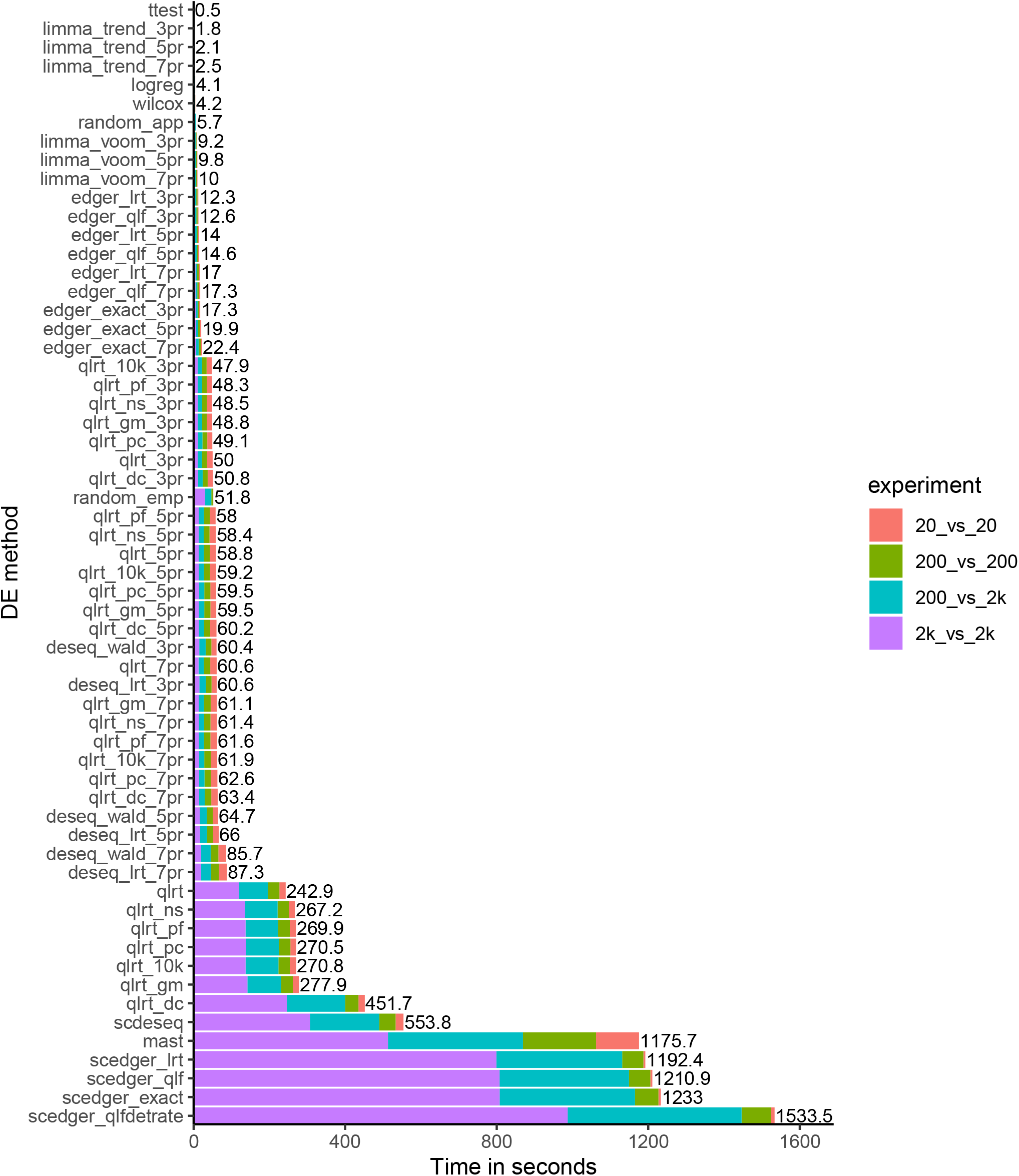
Timing for the different DE methods. For each simulation experiment, the average across replicates is shown. For DE methods that had a runtime dependency on the transformation used, we selected the transformation that resulted in the fastest average runtime.

## References

10x Genomics. 10k Human PBMCs, 3’ v3.1, Chromium Contrller, Cellranger 6.1.0. URL https://www.10xgenomics.com/resources/datasets/10k-human-pbmcs-3-v3-1-chromium-controller-3-1-high.

C. Ahlmann-Eltze and W. Huber. glmGamPoi: Fitting Gamma-Poisson generalized linear models on single cell count data. Bioinformatics, 36(24):5701–5702, 2020. ISSN 14602059. doi: 10.1093/bioinformatics/btaa1009.

C. Ahlmann-Eltze and W. Huber. Comparison of transformations for single-cell RNA-seq data. Nature Methods, page 2021.06.24.449781, apr 2023. ISSN 1548-7091. doi: 10.1038/s41592-023-01814-1. URL https://www.nature.com/articles/s41592-023-01814-1.

S. Anders and W. Huber. Differential expression analysis for sequence count data. Genome Biology, 11(10):R106, oct 2010. ISSN 1474-760X. doi: 10.1186/gb-2010-11-10-r106. URL https://doi.org/10.1186/gb-2010-11-10-r106.

Y. Baran, A. Bercovich, A. Sebe-Pedros, Y. Lubling, A. Giladi, E. Chomsky, Z. Meir, M. Hoichman, A. Lifshitz, and A. Tanay. MetaCell: analysis of single-cell RNA-seq data using K-nn graph partitions. Genome Biology, 20(1):206, dec 2019. ISSN 1474-760X. doi: 10.1186/s13059-019-1812-2. URL https://genomebiology.biomedcentral.com/articles/10.1186/s13059-019-1812-2.

M. Bilous, L. Tran, C. Cianciaruso, A. Gabriel, H. Michel, S. J. Carmona, M. J. Pittet, and D. Gfeller. Metacells untangle large and complex single-cell transcriptome networks. BMC Bioinformatics, 23(1):336, aug 2022. ISSN 1471-2105. doi: 10.1186/s12859-022-04861-1. URL https://doi.org/10.1186/s12859-022-04861-1.

A. S. Booeshaghi, I. B. Hallgrímsdóttir, Á. Gálvez-Merchán, and L. Pachter. Depth normalization for single-cell genomics count data. bioRxiv, page 2022.05.06.490859, 2022. doi: 10.1101/2022.05.06.490859. URL https://www.biorxiv.org/content/10.1101/2022.05.06.490859v1.

W. Chen, Y. Li, J. Easton, D. Finkelstein, G. Wu, and X. Chen. UMI-count modeling and differential expression analysis for single-cell RNA sequencing. Genome Biology, 19(1):70, 2018. ISSN 1474-760X. doi: 10.1186/s13059-018-1438-9. URL https://genomebiology.biomedcentral.com/articles/10.1186/s13059-018-1438-9.

S. Choudhary and R. Satija. Comparison and evaluation of statistical error models for scRNA-seq. Genome Biology, 23(1): 27, 2022. ISSN 1474-760X. doi: 10.1186/s13059-021-02584-9. URL https://doi.org/10.1186/s13059-021-02584-9.

H. L. Crowell, C. Soneson, P. L. Germain, D. Calini, L. Collin, C. Raposo, D. Malhotra, and M. D. Robinson. Muscat Detects Subpopulation-Specific State Transitions From Multi-Sample Multi-Condition Single-Cell Transcriptomics Data. Nature Communications, 11(1):1–12, 2020. ISSN 20411723. doi: 10.1038/s41467-020-19894-4. URL http://dx.doi.org/10.1038/s41467-020-19894-4.

H. L. Crowell, S. X. Morillo Leonardo, C. Soneson, and M. D. Robinson. Built on sand: the shaky foundations of simulating single-cell RNA sequencing data. bioRxiv, 2022. doi: 10.1101/2021.11.15.468676. URL https://www.biorxiv.org/content/early/2022/08/15/2021.11.15.468676.

C. Domínguez Conde, C. Xu, L. B. Jarvis, D. B. Rainbow, S. B. Wells, T. Gomes, S. K. Howlett, O. Suchanek, K. Polanski, H. W. King, L. Mamanova, N. Huang, P. A. Szabo, L. Richardson, L. Bolt, E. S. Fasouli, K. T. Mahbubani, M. Prete, L. Tuck, N. Richoz, Z. K. Tuong, L. Campos, H. S. Mousa, E. J. Needham, S. Pritchard, T. Li, R. Elmentaite, J. Park, E. Rahmani, D. Chen, D. K. Menon, O. A. Bayraktar, L. K. James, K. B. Meyer, N. Yosef, M. R. Clatworthy, P. A. Sims, D. L. Farber, K. Saeb-Parsy, J. L. Jones, and S. A. Teichmann. Cross-tissue immune cell analysis reveals tissue-specific features in humans. Science, 376(6594), may 2022. ISSN 0036-8075. doi: 10.1126/science.abl5197. URL https://www.science.org/doi/10.1126/science.abl5197.

G. Finak, A. McDavid, M. Yajima, J. Deng, V. Gersuk, A. K. Shalek, C. K. Slichter, H. W. Miller, M. J. McElrath, M. Prlic, P. S. Linsley, and R. Gottardo. MAST: a flexible statistical framework for assessing transcriptional changes and characterizing heterogeneity in single-cell RNA sequencing data. Genome Biology, 16(1):278, 2015. ISSN 1474-760X. doi: 10.1186/s13059-015-0844-5. URL http://genomebiology.biomedcentral.com/articles/10.1186/s13059-015-0844-5.

D. Grün, L. Kester, and A. van Oudenaarden. Validation of noise models for single-cell transcriptomics. Nature Methods, 11(6):637–640, 2014. ISSN 1548-7091. doi: 10.1038/nmeth.2930. URL http://www.nature.com/doifinder/10.1038/nmeth.2930.

C. Hafemeister. riffle R package: Non-parametric randomized differential expression test, 2022. URL https://github.com/ChristophH/riffle.

C. Hafemeister and R. Satija. Normalization and variance stabilization of single-cell RNA-seq data using regularized negative binomial regression. Genome Biology, 20(1):296, 2019. ISSN 1474-760X. doi: 10.1186/s13059-019-1874-1. URL https://doi.org/10.1186/s13059-019-1874-1.

Y. Hao, S. Hao, E. Andersen-Nissen, W. M. Mauck, S. Zheng, A. Butler, M. J. Lee, A. J. Wilk, C. Darby, M. Zager, P. Hoffman, M. Stoeckius, E. Papalexi, E. P. Mimitou, J. Jain, A. Srivastava, T. Stuart, L. M. Fleming, B. Yeung, A. J. Rogers, J. M. McElrath, C. A. Blish, R. Gottardo, P. Smibert, and R. Satija. Integrated analysis of multimodal single-cell data. Cell, pages 1–15, 2021. ISSN 00928674. doi: 10.1016/j.cell.2021.04.048. URL https://doi.org/10.1016/j.cell.2021.04.048.

I. Korsunsky, A. Nathan, N. Millard, and S. Raychaudhuri. presto R package: Fast Functions for Differential Expression using Wilcox and AUC, 2020. URL https://github.com/immunogenomics/presto.

J. Lause, P. Berens, and D. Kobak. Analytic Pearson residuals for normalization of single-cell RNA-seq UMI data. Genome Biology, 22(1):258, dec 2021. ISSN 1474-760X. doi: 10.1186/s13059-021-02451-7. URL https://doi.org/10.1101/2020.12.01.405886.

C. W. Law, Y. Chen, W. Shi, and G. K. Smyth. voom: precision weights unlock linear model analysis tools for RNA-seq read counts. Genome Biology, 15(2):R29, 2014. ISSN 1474-760X. doi: 10.1186/gb-2014-15-2-r29. URL https://doi.org/10.1186/gb-2014-15-2-r29.

M. I. Love, W. Huber, and S. Anders. Moderated estimation of fold change and dispersion for RNA-Seq data with DESeq2. Genome Biology, 15(12):550, 2014. ISSN 1465-6906. doi: 10.1186/s13059-014-0550-8. URL http://www.ncbi.nlm.nih.gov/pubmed/25516281.

M. I. Love, S. Anders, and W. Huber. DESeq2 vignette single-cell recommendations, 2022. URL http://bioconductor.org/packages/devel/bioc/vignettes/DESeq2/inst/doc/DESeq2.html#recommendations-for-single-cell-analysis.

A. T. Lun, K. Bach, and J. C. Marioni. Pooling across cells to normalize single-cell RNA sequencing data with many zero counts. Genome Biology, 17(1):1–14, 2016. ISSN 1474760X. doi: 10.1186/s13059-016-0947-7. URL http://dx.doi.org/10.1186/s13059-016-0947-7.

H. B. Mann and D. R. Whitney. On a Test of Whether one of Two Random Variables is Stochastically Larger than the Other. The Annals of Mathematical Statistics, 18(1):50–60, 1947. ISSN 0003-4851. doi: 10.1214/aoms/1177730491.

D. J. McCarthy, Y. Chen, and G. K. Smyth. Differential expression analysis of multifactor RNA-Seq experiments with respect to biological variation. Nucleic Acids Research, 40(10):4288–4297, 2012. doi: 10.1093/nar/gks042. URL http://dx.doi.org/10.1093/nar/gks042.

M. Papadakis, M. Tsagris, M. Dimitriadis, S. Fafalios, I. Tsamardinos, M. Fasiolo, G. Borboudakis, J. Burkardt, C. Zou, K. Lakiotaki, and C. Chatzipantsiou. Rfast: A Collection of Efficient and Extremely Fast R Functions, 2022. URL https://github.com/RfastOfficial/Rfast.

S. Persad, Z.-N. Choo, C. Dien, N. Sohail, I. Masilionis, R. Chaligné, T. Nawy, C. C. Brown, R. Sharma, I. Pe’er, M. Setty, and D. Pe’er. SEACells infers transcriptional and epigenomic cellular states from single-cell genomics data. Nature Biotechnology, page 2022.04.02.486748, mar 2023. ISSN 1087-0156. doi: 10.1038/s41587-023-01716-9. URL https://www.nature.com/articles/s41587-023-01716-9.

J. M. Pullin and D. J. McCarthy. A comparison of marker gene selection methods for single-cell RNA sequencing data. bioRxiv, 2022. doi: 10.1101/2022.05.09.490241. URL https://www.biorxiv.org/content/early/2022/09/14/2022.05.09.490241.

R Core Team. R: A Language and Environment for Statistical Computing. R Foundation for Statistical Computing, Vienna, Austria, 2021. URL https://www.r-project.org/.

M. E. Ritchie, B. Phipson, D. Wu, Y. Hu, C. W. Law, W. Shi, and G. K. Smyth. limma powers differential expression analyses for RNA-sequencing and microarray studies. Nucleic Acids Research, 43(7):e47–e47, apr 2015. ISSN 1362-4962. doi: 10.1093/nar/gkv007. URL http://academic.oup.com/nar/article/43/7/e47/2414268/limma-powers-differential-expression-analyses-for.

R. Satija, J. a. Farrell, D. Gennert, A. F. Schier, and A. Regev. Spatial reconstruction of single-cell gene expression data. Nature Biotechnology, 33(5), 2015. ISSN 1087-0156. doi: 10.1038/nbt.3192. URL http://www.nature.com/doifinder/10.1038/nbt.3192.

C. Soneson and M. D. Robinson. Bias, robustness and scalability in single-cell differential expression analysis. Nature Methods, 15(4):255–261, 2018. ISSN 15487105. doi: 10.1038/nmeth.4612. URL http://dx.doi.org/10.1038/nmeth.4612.

J. W. Squair, M. Gautier, C. Kathe, M. A. Anderson, N. D. James, T. H. Hutson, R. Hudelle, T. Qaiser, K. J. E. Matson, Q. Barraud, A. J. Levine, G. La Manno, M. A. Skinnider, and G. Courtine. Confronting false discoveries in single-cell differential expression. Nature Communications, 12(1):5692, 2021. ISSN 2041-1723. doi: 10.1038/s41467-021-25960-2. URL https://doi.org/10.1038/s41467-021-25960-2.

B. L. Welch. THE GENERALIZATION OF ‘STUDENT’S’ PROBLEM WHEN SEVERAL DIFFERENT POPULATION VARLANCES ARE INVOLVED. Biometrika, 34(1-2):28–35, 1947. ISSN 0006-3444. doi: 10.1093/biomet/34.1-2.28. URL https://doi.org/10.1093/biomet/34.1-2.28.

F. A. Wolf, P. Angerer, and F. J. Theis. SCANPY: Large-scale single-cell gene expression data analysis. Genome Biology, 19(1):1–5, 2018. ISSN 1474760X. doi: 10.1186/s13059-017-1382-0.

