## Supplementary figures and images for "Single-cell RNA-seq differential expression tests within a sample should use pseudo-bulk data of pseudo-replicates"

### Additional File 2

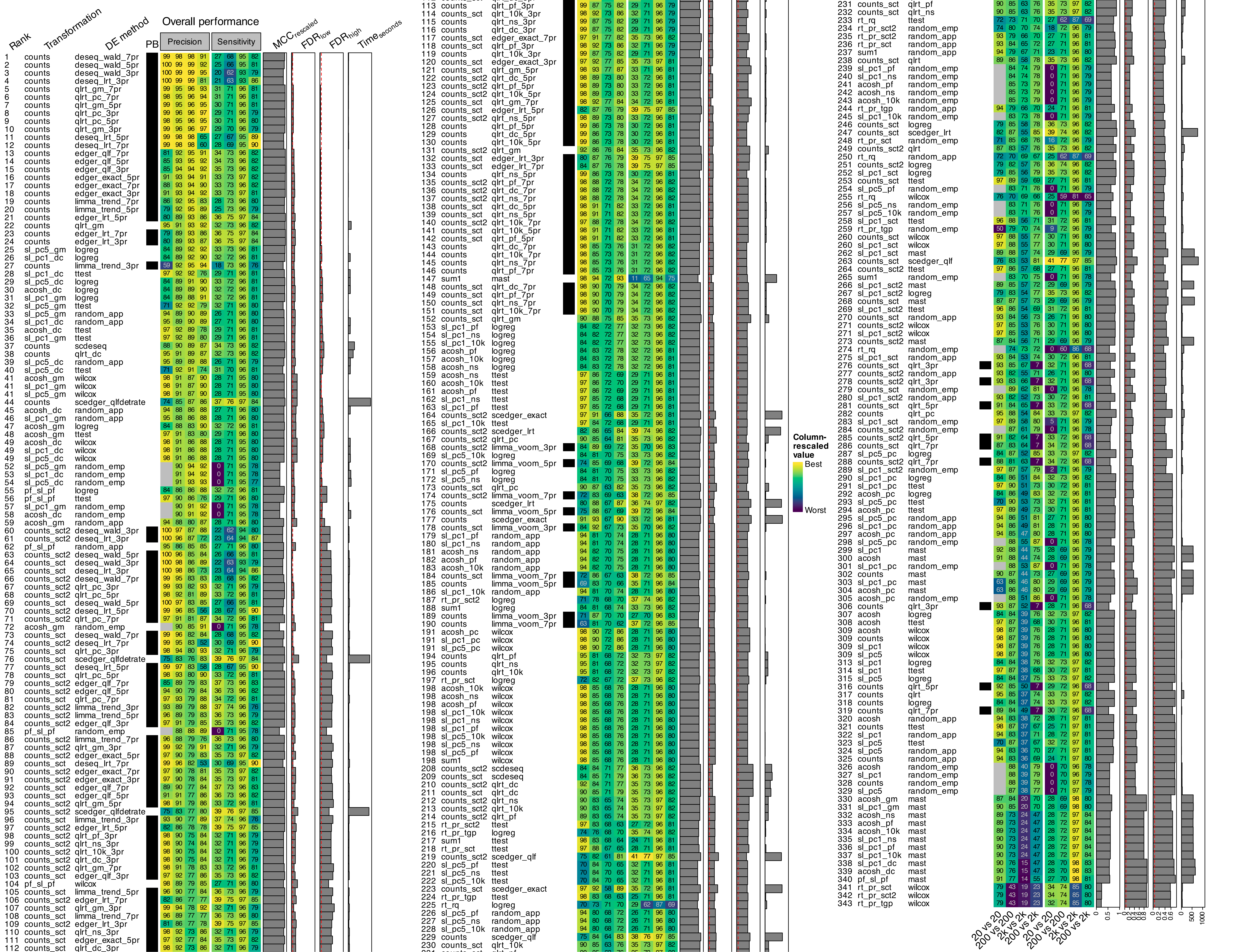

### Additional File 4

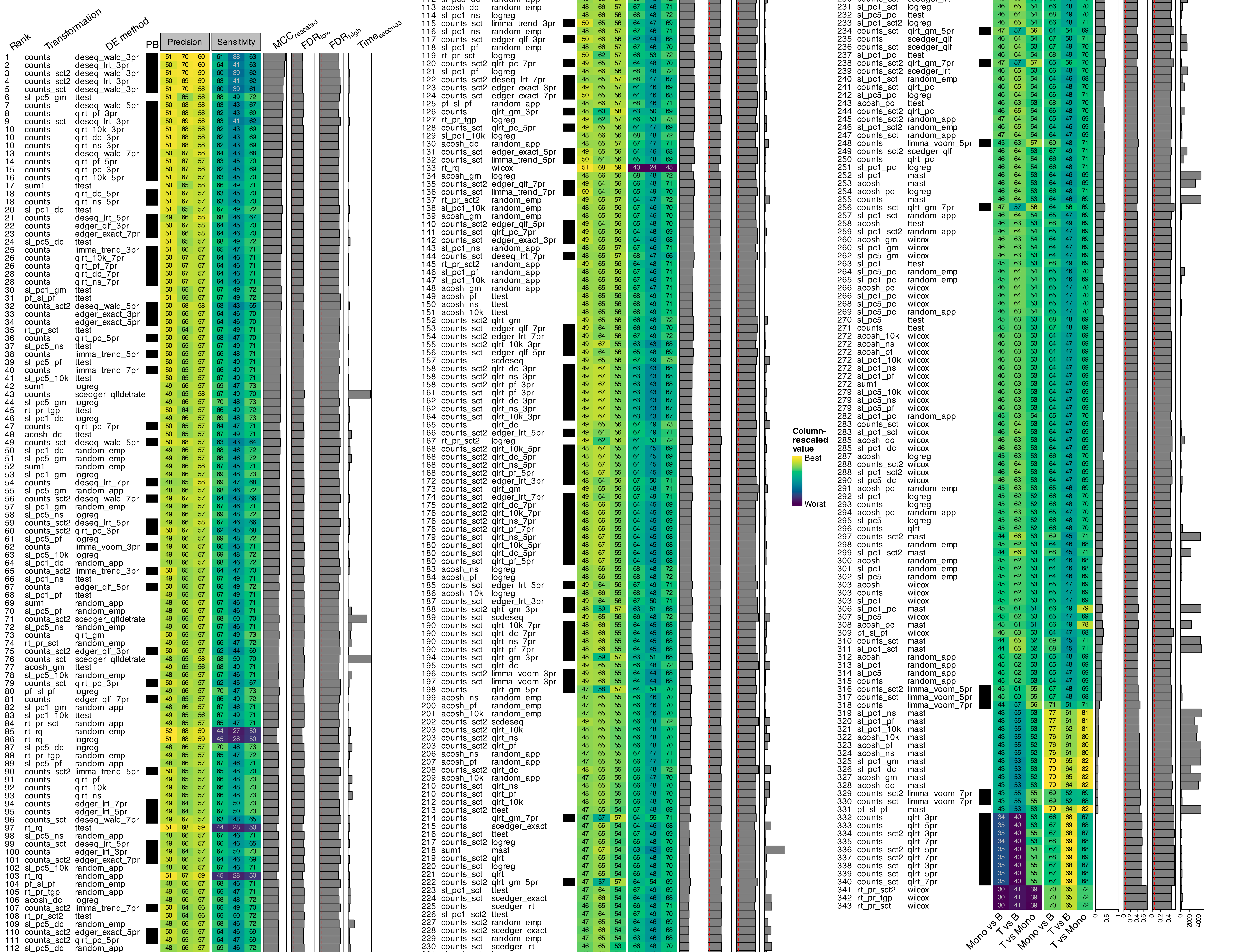
